## Supplemental Information for "Gene regulatory dynamics during craniofacial development in a carnivorous marsupial"

### Supplementary Figures

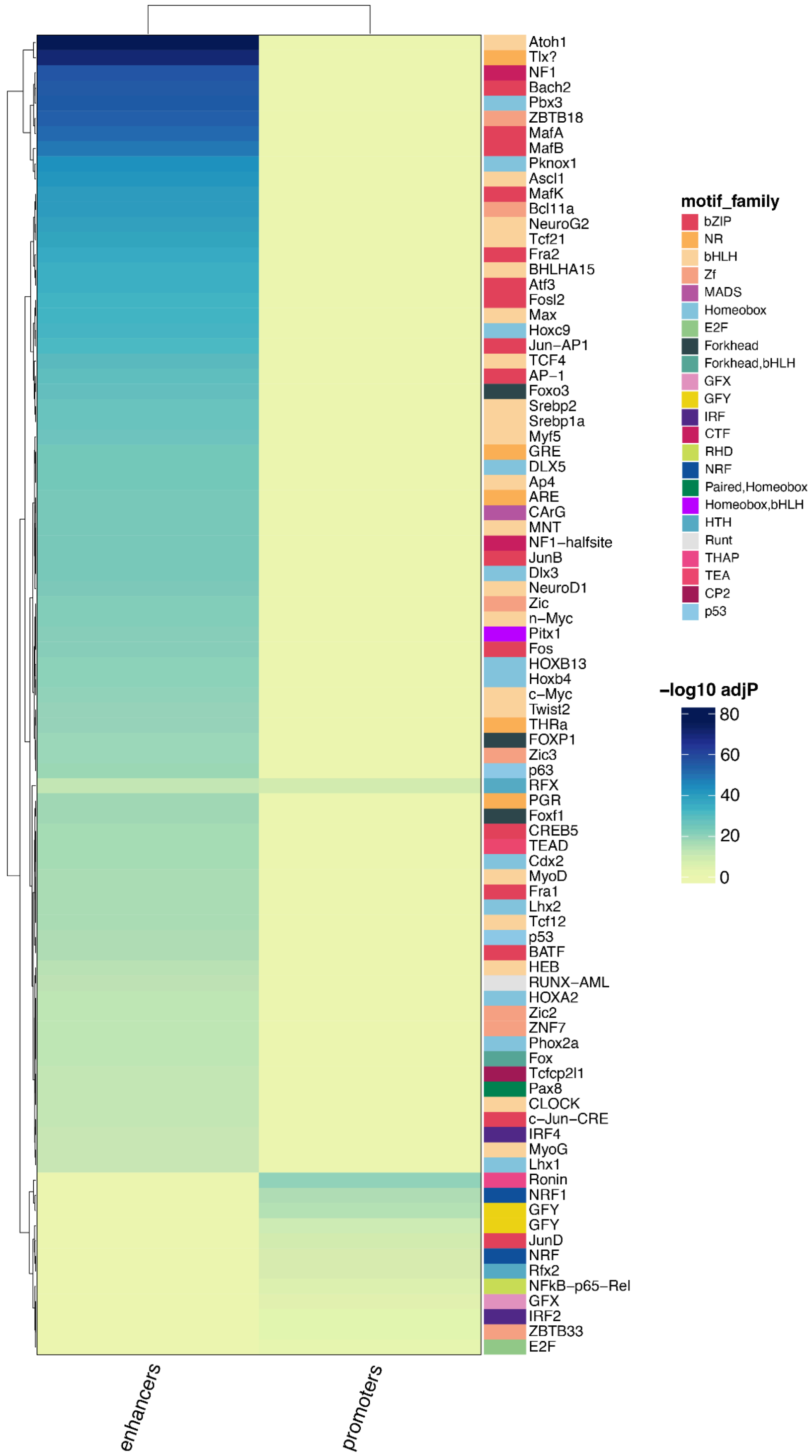


**Supplementary Figure 1. Homer motif enrichment for dunnart promoters and enhancers for the top 20 enriched TF families.** $\text{Log}_{10}$ FDR adjusted p-value scores are shown as a heatmap in enhancers and promoters. Individual TFs are labeled on the right hand side of the heatmap. Distinct colors represent TF families.

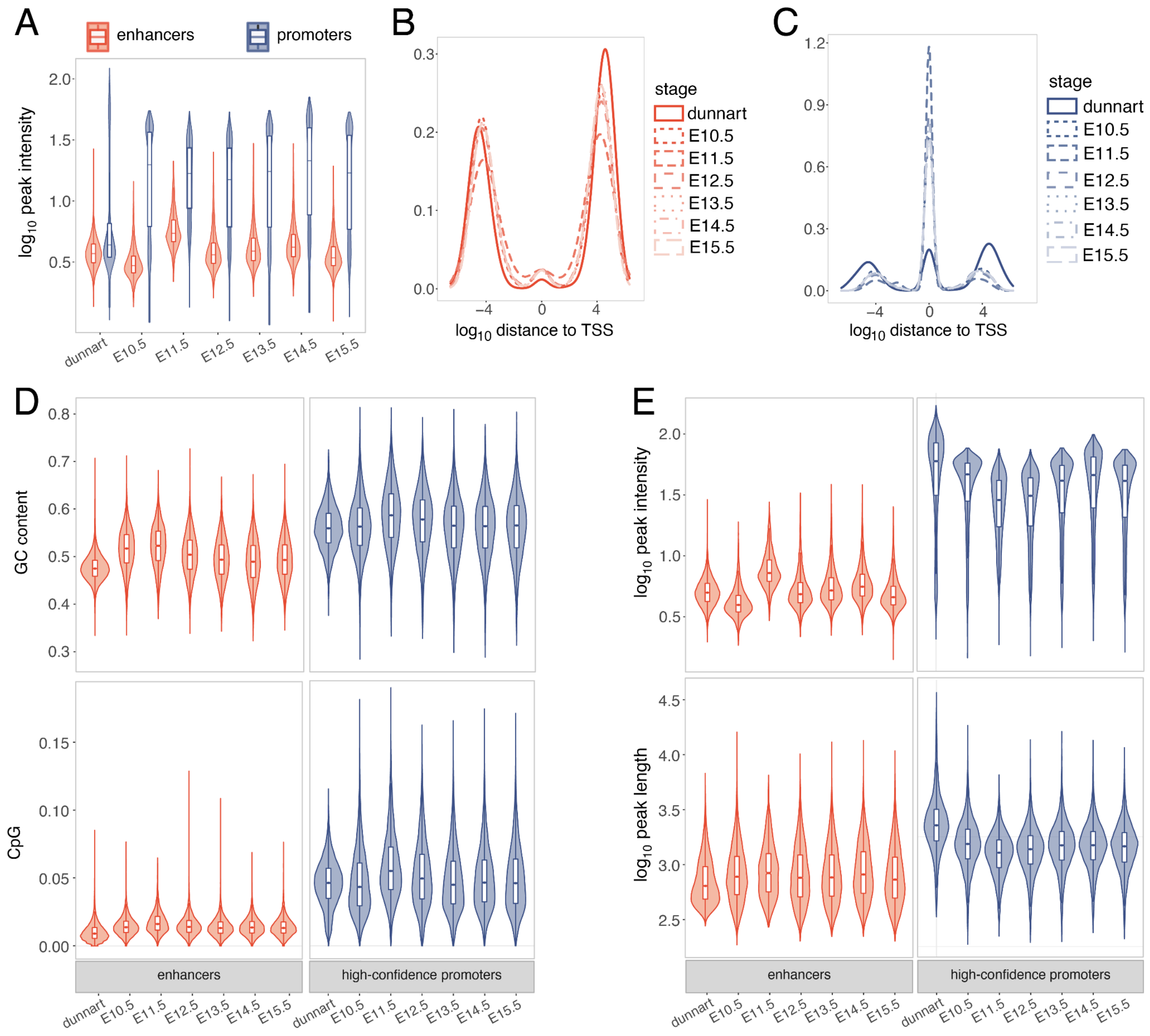


**Supplementary Figure 2. Features of predicted promoters and enhancers in the dunnart and mouse.** (A) $Log_{10}$ peak intensity (measure of enrichment) for enhancers (orange) and promoters (blue) prior to filtering. (B) $Log_{10}$ distance to the nearest TSS for enhancers (orange) and promoters (blue). After clustering and filtering for high-confidence promoters we observed consistent patterns for features including, (D) CpG and GC content and, (E) $Log_{10}$ peak intensity and $Log_{10}$ peak length.

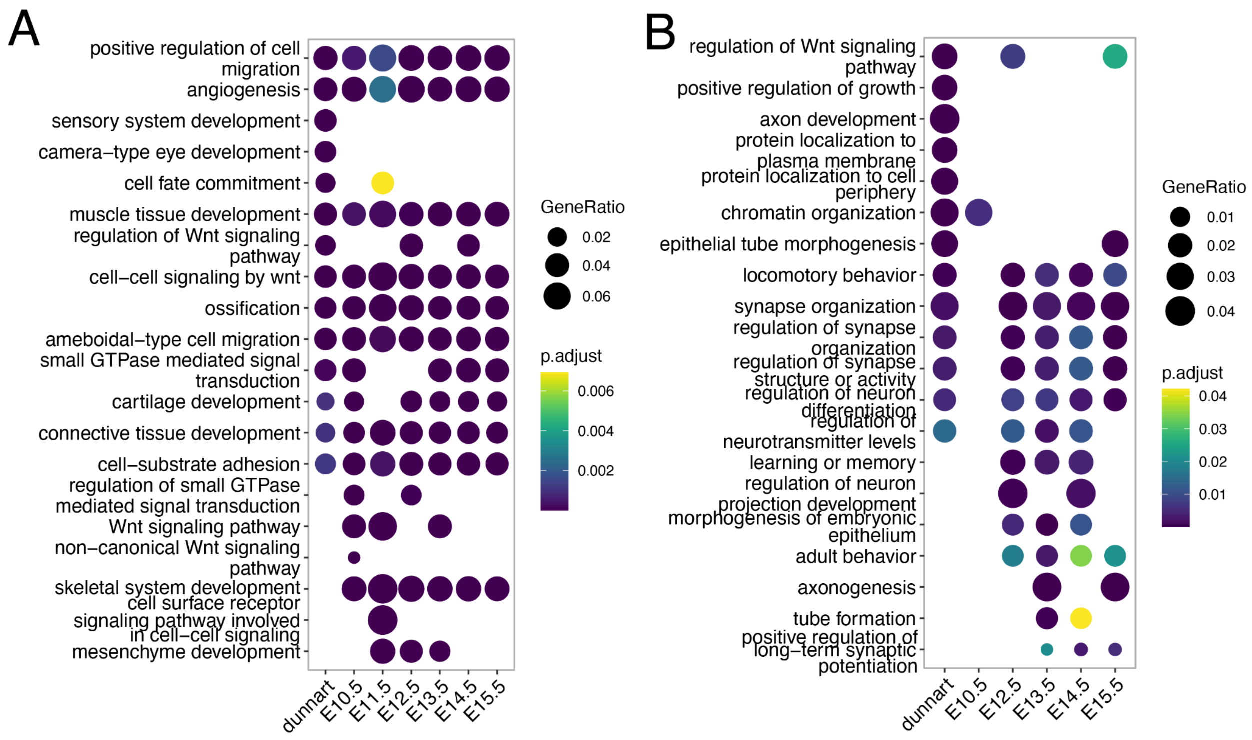


**Supplementary Figure 3. Top GO enriched terms in the dunnart and mouse embryonic stages** for (A) genes near enhancers and, (B) genes near high-confidence promoters.

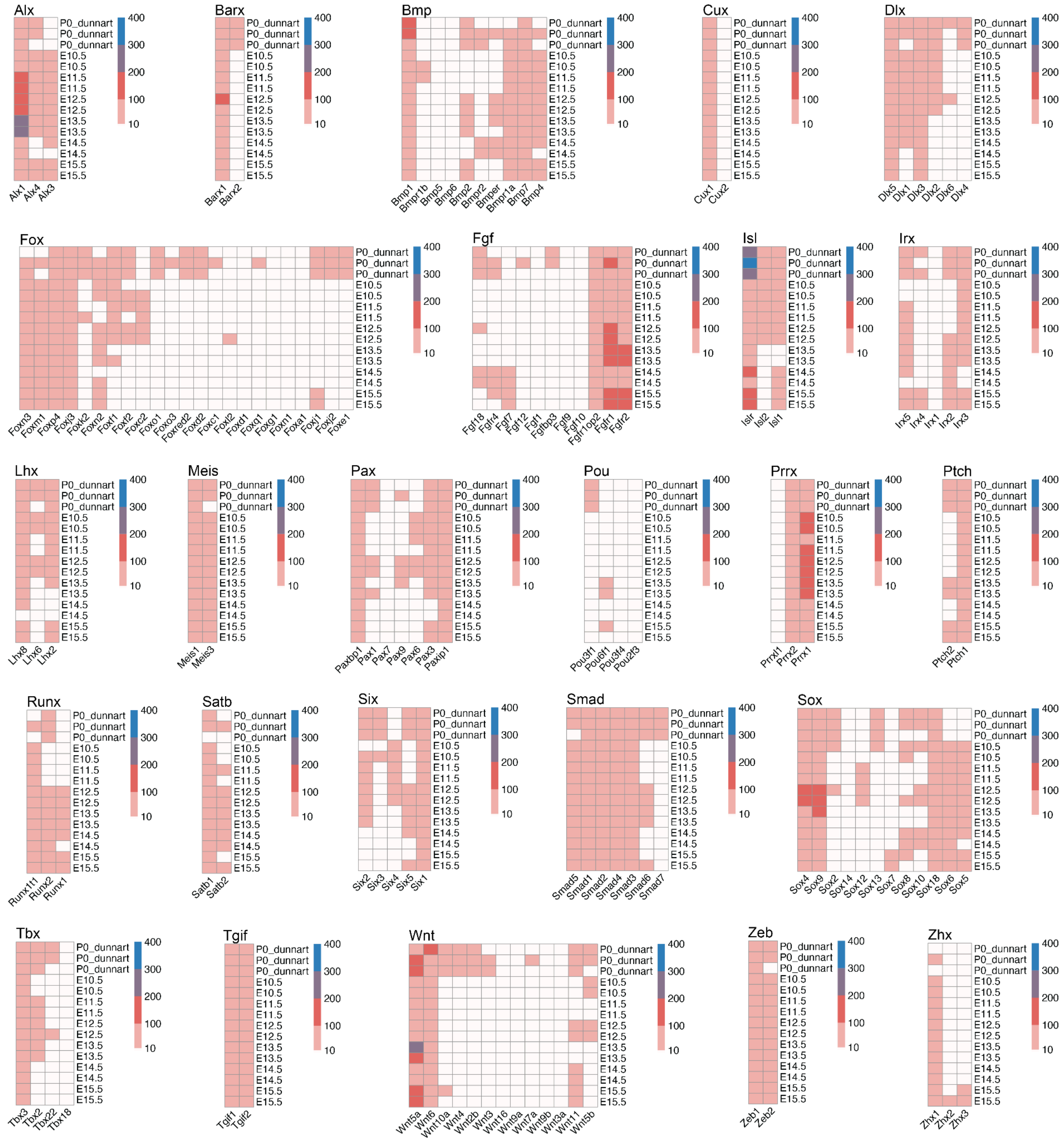


**Supplementary Figure 4. Heatmaps showing gene expression values (TPM) for common developmental genes across dunnart and mouse.**

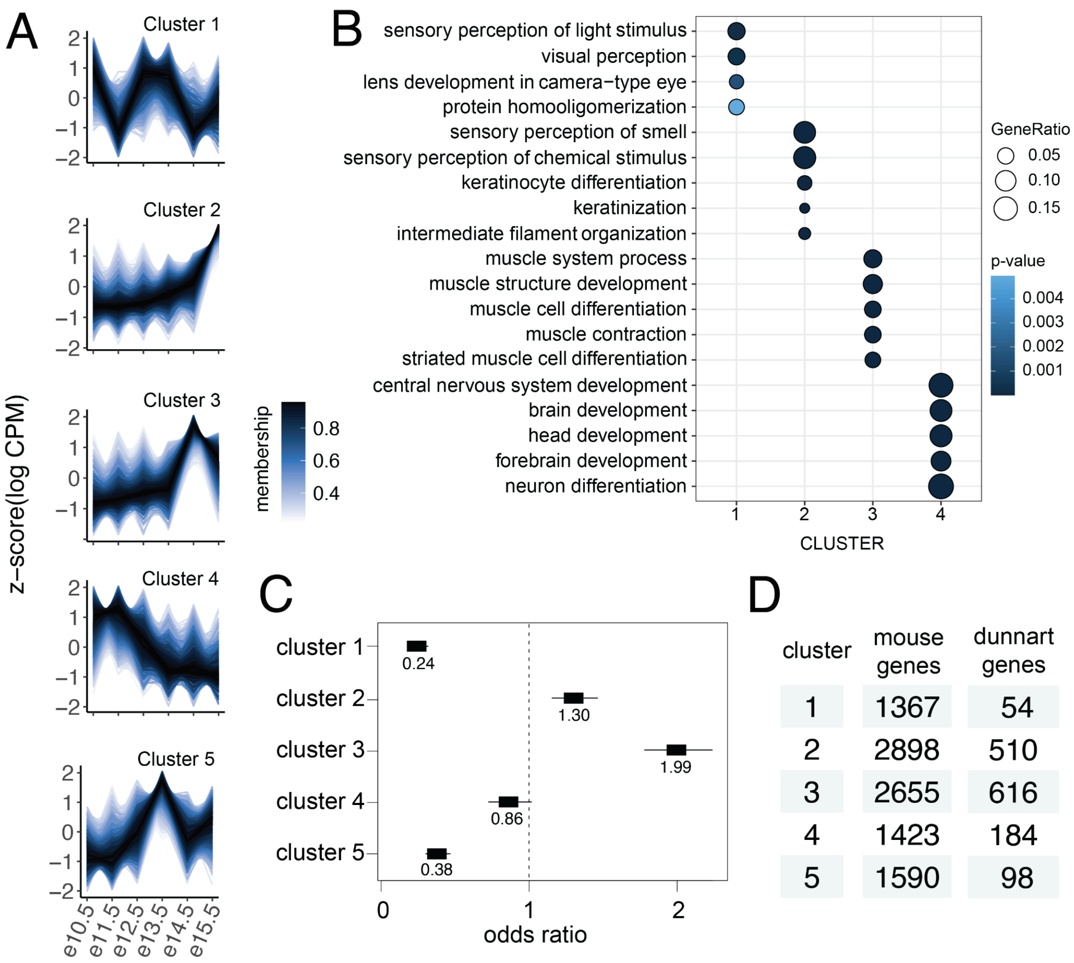


| cluster | mouse genes | dunnart genes |
|---|---|---|
| 1 | 1367 | 54 |
| 2 | 2898 | 510 |
| 3 | 2655 | 616 |
| 4 | 1423 | 184 |
| 5 | 1590 | 98 |

**Supplementary Figure 5. Temporal gene expression dynamics.** (A) Z-scaled temporal expression (logCPM) plotted across embryonic timepoints for five clusters. Membership values are used to indicate to what degree a data point belongs to each cluster. (B) Top enriched biological processes for each cluster (FDR corrected p<0.01, background gene set is all differentially expressed genes used for clustering). (C) Odds ratio estimates with 95% confidence interval for genes expressed in dunnart in each cluster. (D) Number of genes in each cluster for mouse and dunnart.

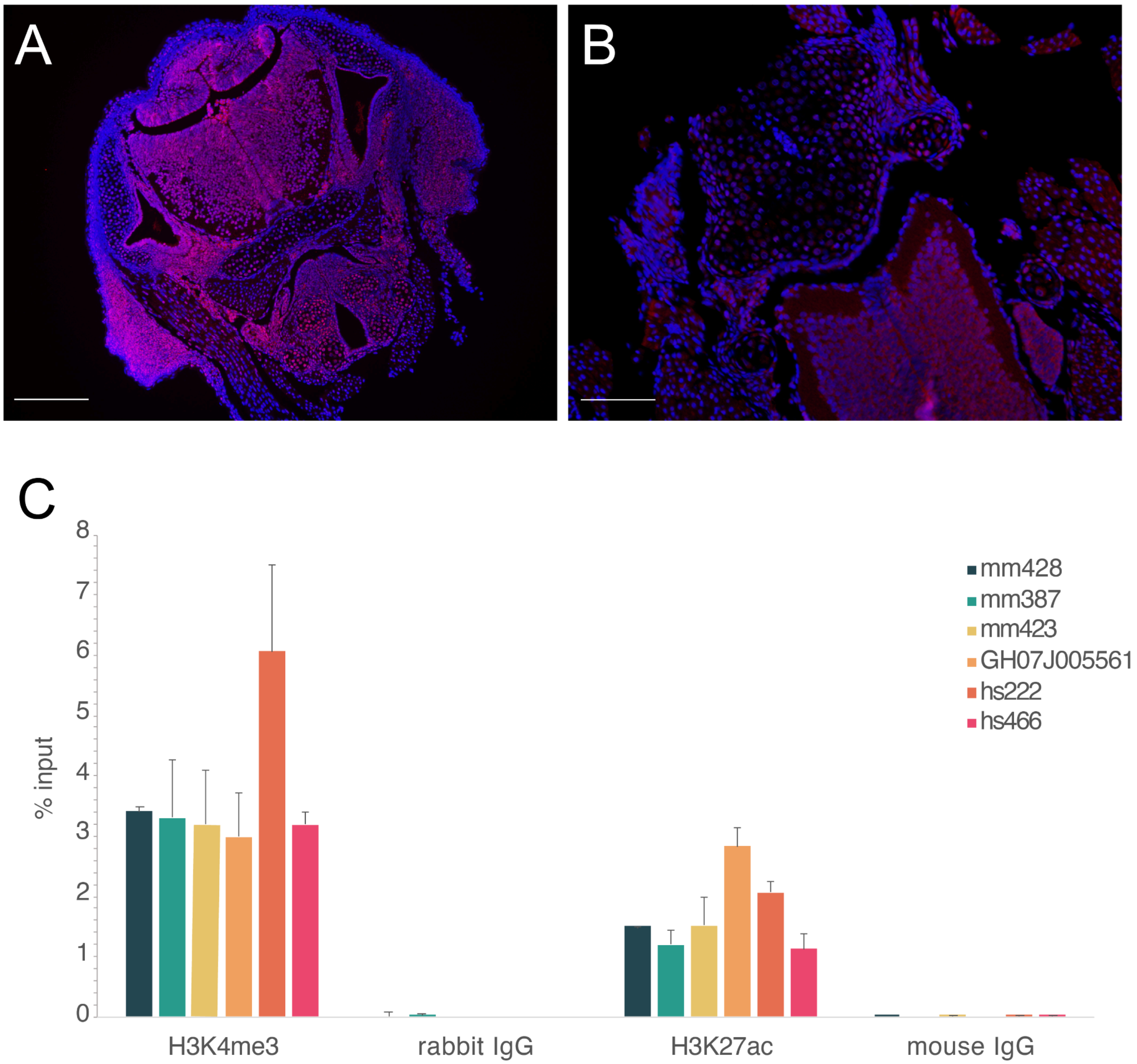


**Supplementary Figure 6. Validation of antibodies in dunnart craniofacial tissue with immunofluorescence and qPCR.** Localisation of (A) H3K27ac (pink, scale bar = 250um) and (B) H3K4me3 (pink, scale bar = 80um), in D0 dunnart head sections with nuclei stained with DAPI (blue) (C) enhancer regions expected to be enriched in the IP samples presented as the percentage of the input control sample as measured by qPCR.

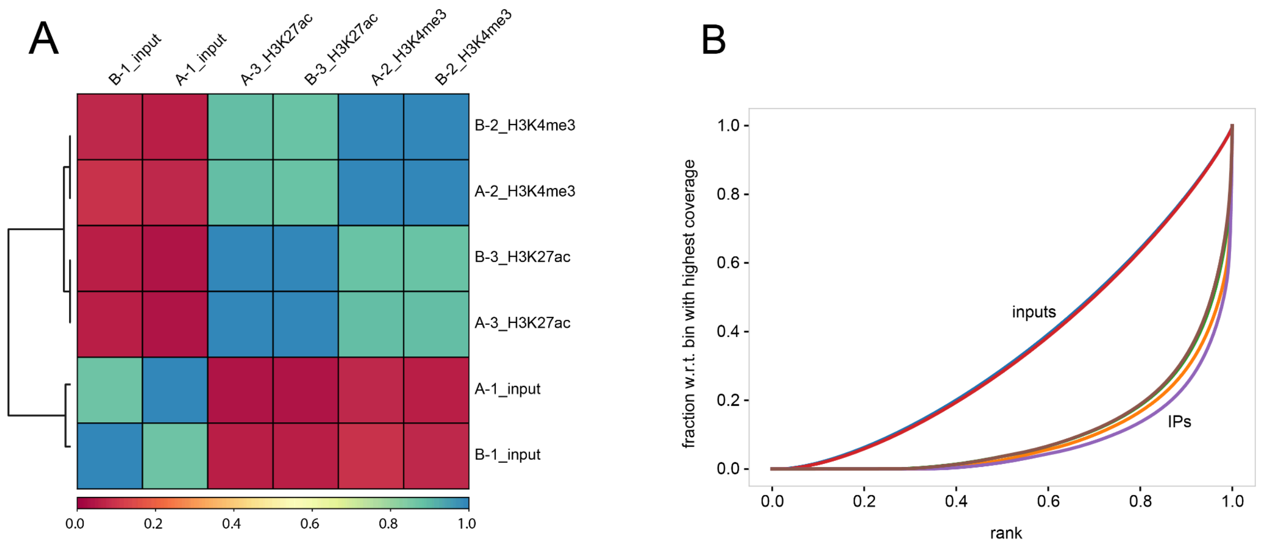


**Supplementary Figure 7. deepTools quality control plots for dunnart subsampled aligned BAM files.** (A). Overall similarity between BAM files based on read coverage within genomic regions with Pearson correlation coefficients plotted for H3K27ac, H3K4me3 and input control. (B) Fingerprint plot showing a profile of cumulative read coverages for each BAM file. All reads overlapping a window (bin) of the specified length are counted, sorted and plotted for H3K27ac, H3K4me3 and input control.

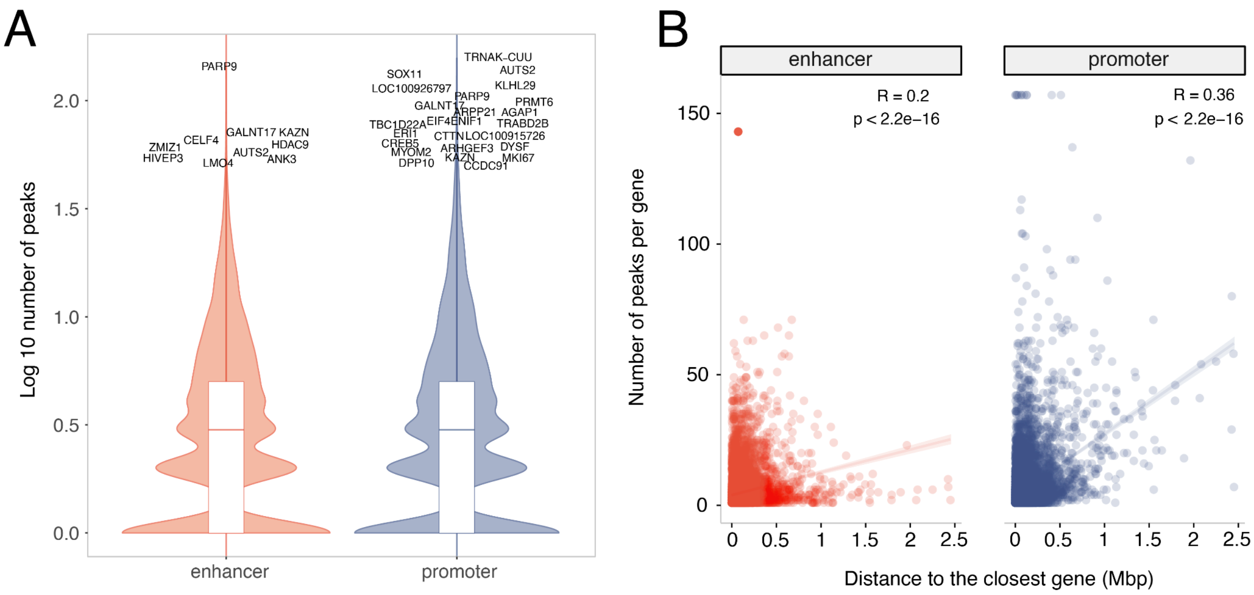


**Supplementary Figure 8. Number of peaks per gene for enhancer- and promoter- associated peaks.** (A) $Log_{10}$ number of peaks per gene. Genes with greater than 50 peaks are noted. (B) Scatter-plot with distance to the closest gene on the x-axis and number of peaks per gene on the y-axis. There is a weak but significant correlation between the number of peaks per gene and the distance to the next closest gene in both enhancers and promoters.

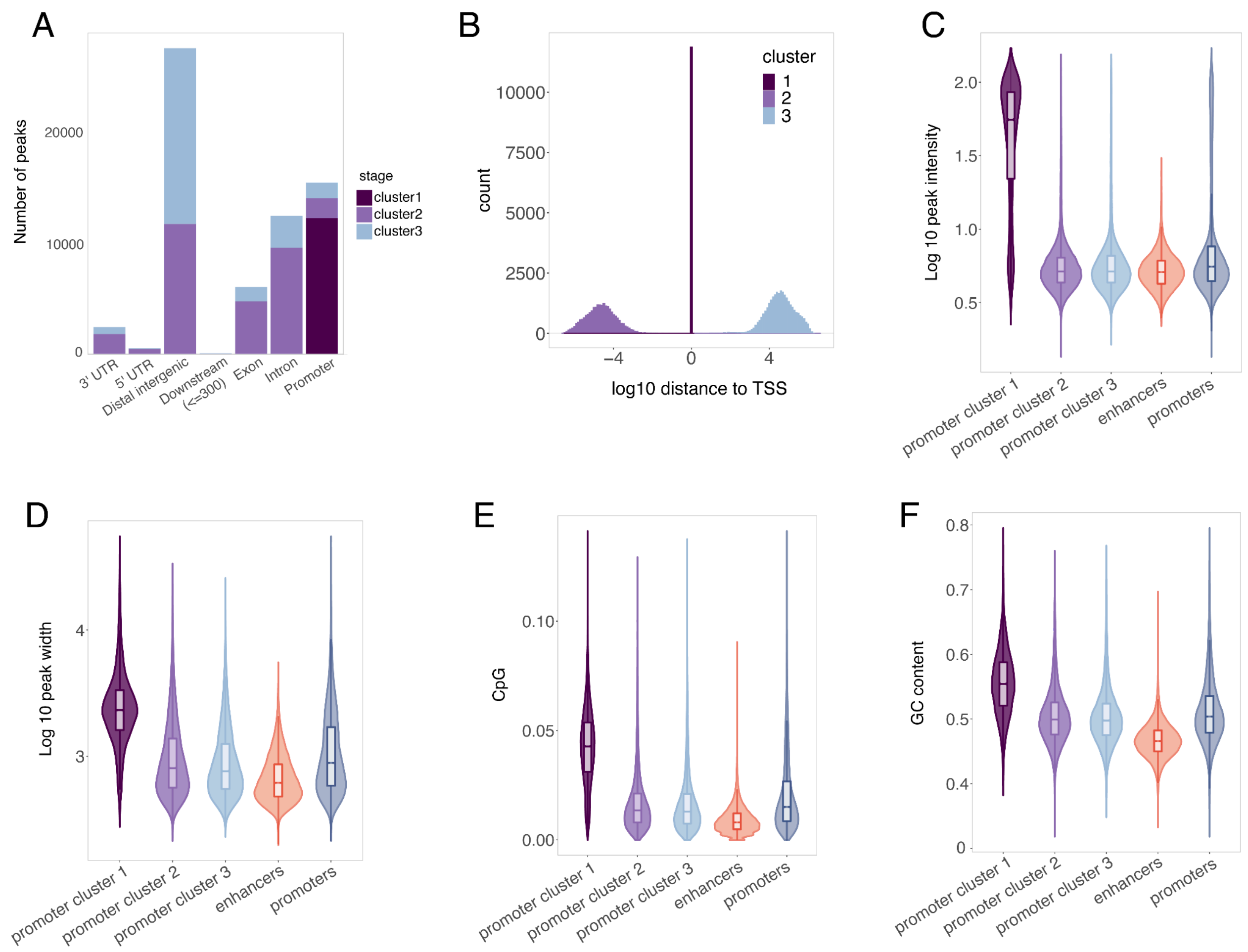


**Supplementary Figure 9. Subsetting high-confidence promoter-associated peaks with k-means clustering.** (A) Barplot for the number of promoter-associated peaks per cluster and histogram showing the distribution of peak distance from the nearest TSS in each cluster. (B) Genomic annotations for promoter-associated peaks in each cluster. (C) GC content, (D) $Log_{10}$ peak length, (E) CpG content, and (F) $Log_{10}$ of clustered promoter-associated peaks, enhancer-associated peaks, and unclustered promoter-associated peaks. Statistical significance (Wilcoxon, FDR-adjusted, $p <$ 0.00001) compared to cluster 1 promoter-associated peaks is denoted by ****.

A

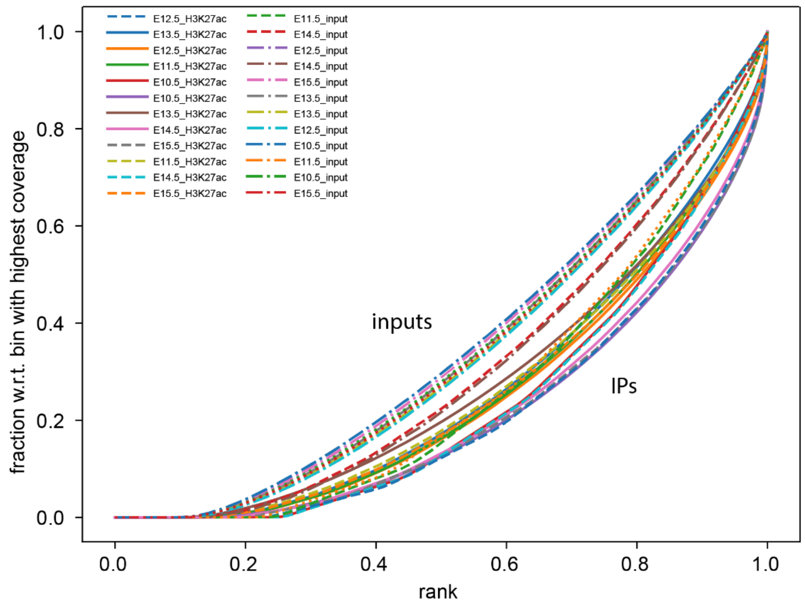


B

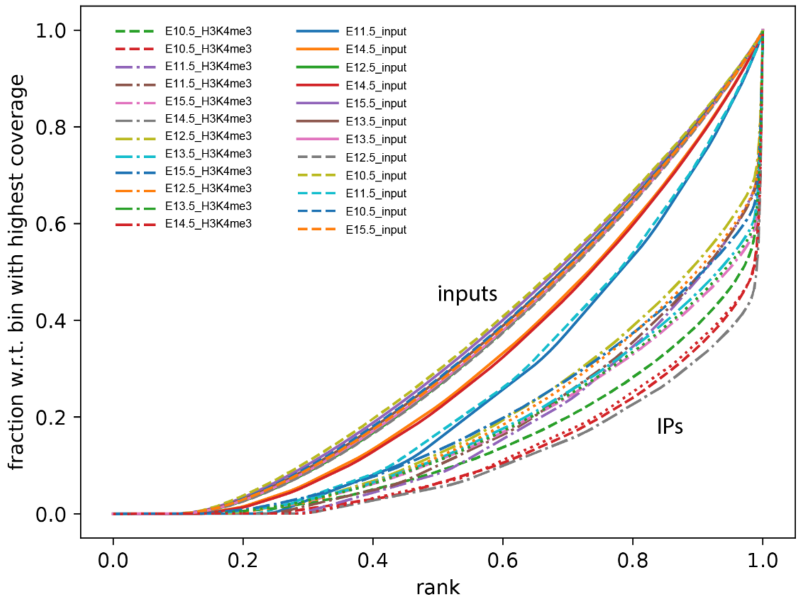


**Supplementary Figure 10. deepTools fingerprint plot for mouse subsampled aligned BAM files.** Cumulative read coverages for each BAM file. All reads overlapping a window (bin) of the specified length are counted, sorted and plotted for (A) H3K27ac, (B) H3K4me3.

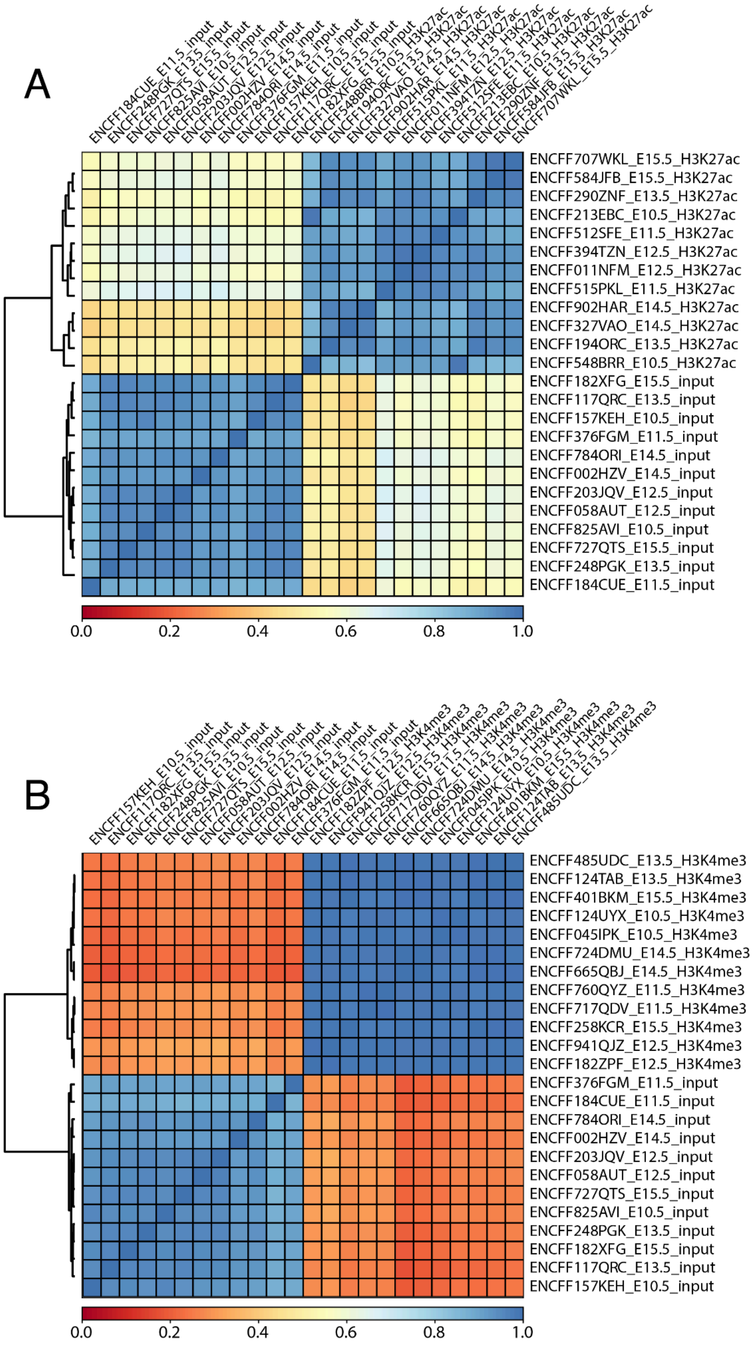


**Supplementary Figure 11. deepTools correlation plots for mouse subsampled aligned BAM files.** Overall similarity between BAM files based on read coverage within genomic regions with Pearson correlation coefficients plotted for (A) H3K27ac, (B) H3K4me3

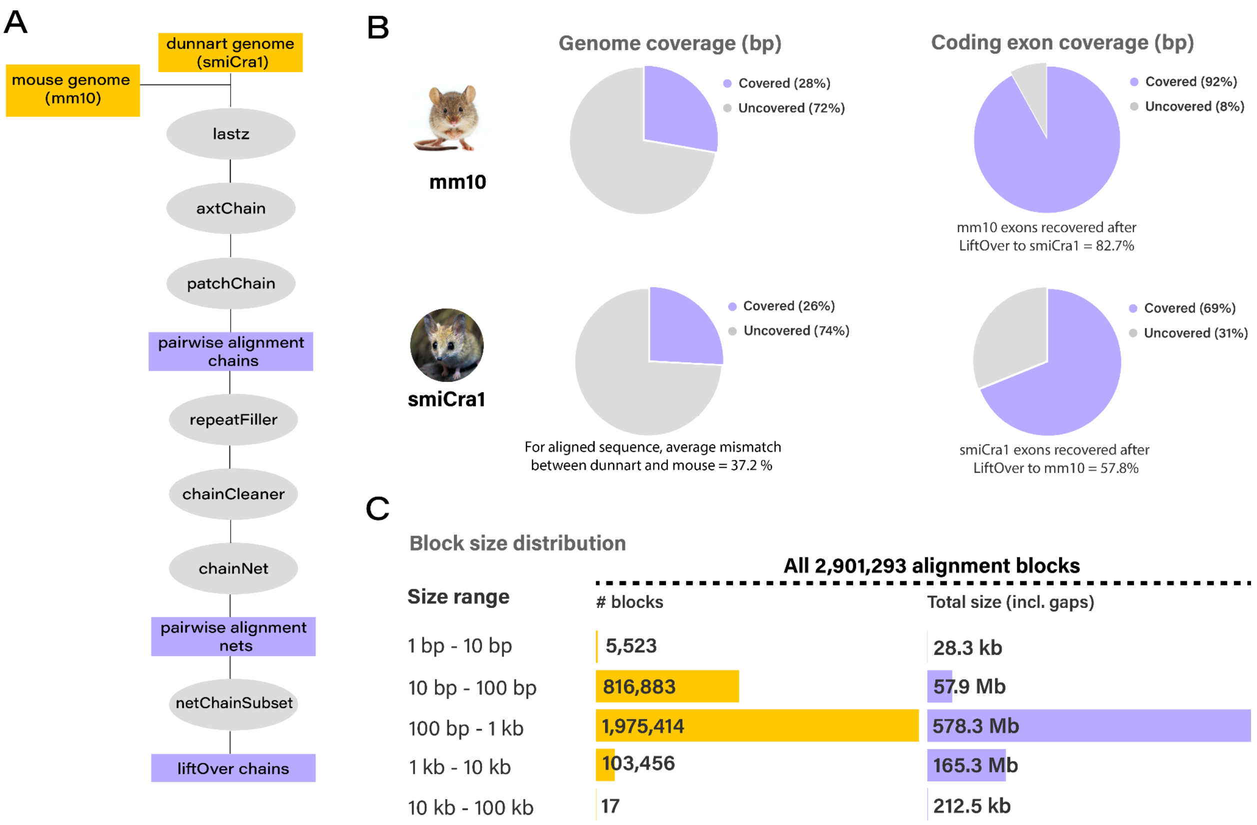


**Supplementary Figure 12. Genome alignment between the fat-tailed dunnart (*Sminthopsis crassicaudata* and mouse (*Mus musculus)*.** (A) Genome alignment workflow. (B) Genome coverage and exon coverage (bp) between mouse and dunnart. Coverage of exons recovered after LiftOver. (C) Block size distribution for the dunnart including number of blocks and total size of the blocks.

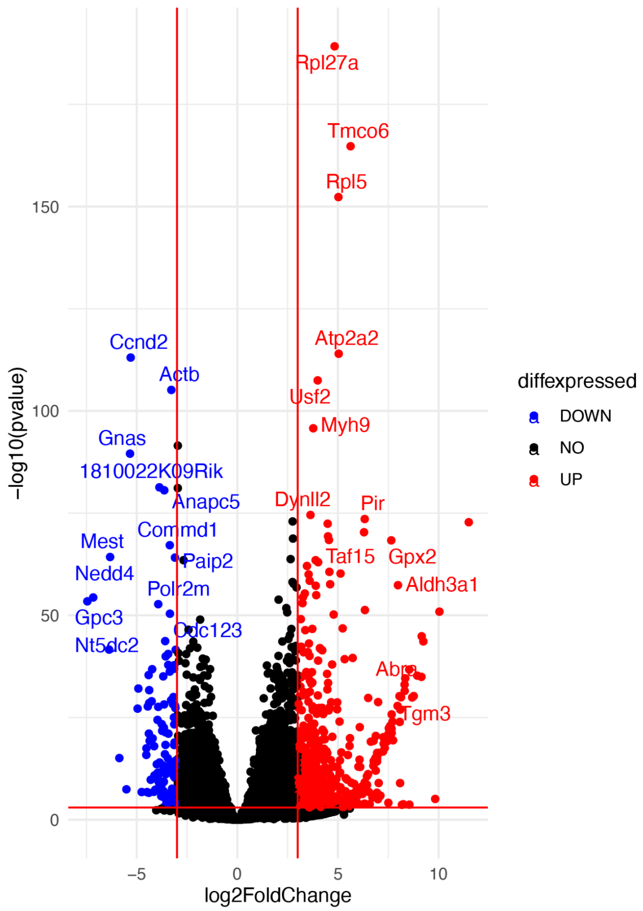


**Supplementary Figure 13. Volcano plot for both upregulated and downregulated differentially expressed genes from mouse (any stage) and dunnart.** Stringent log2 fold change and pvalue cutoffs were used given the comparisons between datasets from two different species.
